# GeneQC: A quality control tool for gene expression estimation based on RNA-sequencing reads mapping

**DOI:** 10.1101/266445

**Authors:** Adam McDermaid, Xin Chen, Yiran Zhang, Juan Xie, Cankun Wang, Qin Ma

## Abstract

**Motivation:** One of the main benefits of using modern RNA-sequencing (RNA-Seq) technology is the more accurate gene expression estimations compared with previous generations of expression data, such as the microarray. However, numerous issues can result in the possibility that an RNA-Seq read can be mapped to multiple locations on the reference genome with the same alignment scores, which occurs in plant, animal, and metagenome samples. Such a read is so-called a multiple-mapping read (MMR). The impact of these MMRs is reflected in gene expression estimation and all downstream analyses, including differential gene expression, functional enrichment, etc. Current analysis pipelines lack the tools to effectively test the reliability of gene expression estimations, thus are incapable of ensuring the validity of all downstream analyses.

**Results:** Our investigation into 95 RNA-Seq datasets from seven species (totaling 1,951GB) indicates an average of roughly 22% of all reads are MMRs for plant and animal species. Here we present a tool called ***GeneQC*** (**Gene** expression **Q**uality **C**ontrol), which can accurately estimate the reliability of each gene’s expression level. The underlying algorithm is designed based on extracted genomic and transcriptomic features, which are then combined using elastic-net regularization and mixture model fitting to provide a clearer picture of mapping uncertainty for each gene. GeneQC allows researchers to determine reliable expression estimations and conduct further analysis on the gene expression that is of sufficient quality. This tool also enables researchers to investigate continued re-alignment methods to determine more accurate gene expression estimates for those with low reliability.

**Availability:** GeneQC is freely available at http://bmbl.sdstate.edu/GeneQC/home.html.

**Supplementary information:** Supplementary data are available at *Bioinformatics* online.

## 1 Introduction

RNA-Seq is a revolutionary high-throughput process that allows researchers to observe the genetic makeup of a particular sample (Garber, et al., 2011; Ozsolak and Milos, 2011; Wang, et al., 2009). Research involving RNA-Seq data produces genetic expression profiles, in which a discrete expression value for each annotated gene for that species is identified. These gene expression profiles are extracted through computational RNA-Seq analysis pipelines (Anders, et al., 2015; Andrews, 2010; Bonfert, et al., 2015; Chang, et al., 2015; Dobin, et al., 2013; Grabherr, et al., 2011; Kim, et al., 2015; Kong, 2011; Li and Dewey, 2011; Pertea, et al., 2016; Pertea, et al., 2015; Philippe, et al., 2013; Trapnell, et al., 2009; Wang, et al., 2010; Wang, et al., 2009; Wu, et al., 2013; Wu, et al., 2016; Yuan, et al., 2017), which can be analyzed further to identify differentially expressed genes between treatment groups (Anders and Huber, 2012; Pimentel, et al., 2017; Ritchie, et al., 2015; Robinson, et al., 2010; Trapnell, et al., 2012), enriched functional gene modules (Chen, et al., 2009; Pathan, et al., 2015; Subramanian, et al., 2005; Zhou and Su, 2007), co-expression networks (Li, et al., 2009), and to generate visualizations to assist in broad interpretations between treatment groups (Ge, 2017; Goff, et al., 2013; Harshbarger, et al., 2017; McDermaid, et al., 2018; Nelson, et al., 2017; Nueda, et al., 2017; Perkel, 2018; Powell, 2015; Younesy, et al., 2015), among other applications.

One of the applications of RNA-Seq analysis pipelines is to use the sequenced short reads with a reference genome, if available, to estimate the expression level of each gene (Miller, et al., 2014; Nagalakshmi, et al., 2008). The basic process is to map these short reads to the location with the best alignment score on the reference genome (Wu, et al., 2014). Even though numerous methods have been developed to facilitate the analysis of RNA-Seq data, some critical issues persist. The nature of DNA—long strands of millions of base-pairs created by a reordering of the four nucleotides—makes it inevitable that some similarities and duplications will occur throughout the genome. This can lead to ambiguity during read mapping, with specific reads being aligned to multiple locations across the reference genome with the same alignment scores (Baruzzo, et al., 2017; D’haeseleer, 2006; Li, et al., 2009; Oshlack, et al., 2010; Swan, 2013; Trapnell, et al., 2013).

This MMR problem can be observed in any genomic region, including, exons and transcripts. For conciseness, we refer to these genomic regions simply as “genes.” This issue has been observed in many diploid species, including human and other mammals and Arabidopsis (Anders and Huber, 2012; Anders, et al., 2015; Bonfert, et al., 2015; Garber, et al., 2011; Wang, et al., 2009), as well as many multiploid species.

The general solution of the MMR problem in previous studies is to discard or evenly distribute to all potential locations, leading to severe, biased underestimation or overestimation of the gene expression levels, respectively (Chang, et al., 2015). More commonly, a proportional assignment of ambiguous reads, in which the read is segmented in smaller portions based on the number of possible mapping locations and uniquely mapped reads to each of them (Li, et al., 2009). In species with high levels of uncertainty, especially angiosperms, the MMR problem can have serious implications on gene expression levels and can be extremely hard to remediate due to the genes’ and chromosomes’ duplicative nature (Grabherr, et al., 2011). In some species, such as Glycine, up to 75% of the genes have the duplicated partners in its genome (Kim, et al., 2015). During our initial investigation into the MMR problem, 95 datasets totaling 1,951GB were analyzed, and it was determined that an average of 22% of all reads were ambiguously aligned over seven distinct plant and animal species (Figure 1A). In some datasets, over 35% of the reads were ambiguously aligned, with more details are provided in *Preliminary Analysis S1* and *Table S1*.

**Figure 1.**
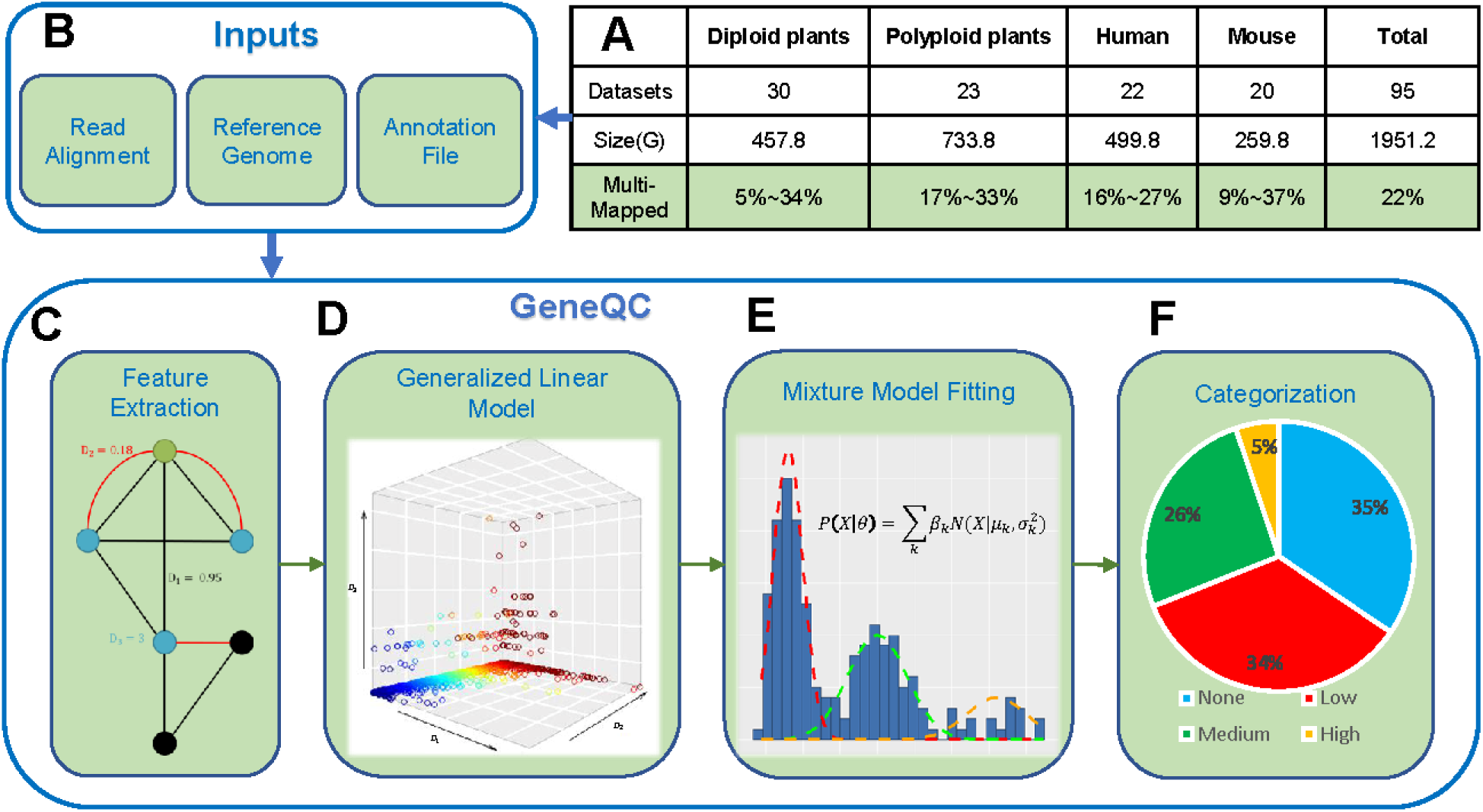
*(A)* The MMR percentages for the 95 analyzed datasets across seven species. More detailed information can be found in Table S1; (B) GeneQC takes a read alignment, reference genome, and annotation file as inputs; (C) The first step of GeneQC is to extract features related to mapping uncertainty for each annotated gene; (D) Using the extracted features, elastic-net regularization is used to calculate the D-score, which represents the mapping uncertainty for each gene; (E) A series of Mixture Normal and Mixture Gamma distributions are fit to the D-scores; (F) The mixture models are used to categorize the D-scores into different levels of mapping uncertainty along with a statistical alternative likelihood value for each gene.

To address this issue, we present GeneQC based on novel applications of robust regression and mixture model fitting approaches to quantify the mapping uncertainty issue. This tool can determine the genes having reliable expression estimates and those require further analysis, along with a statistical significant evaluation of the mapping uncertainty level. The basic idea is to develop a distinct score, referred to as D-score, to group genes into several categorizations with different reliability levels, through integration and modeling of genomic and transcriptomic features. Specifically, (i) sequence similarity between a particular gene and other genes is collected to give an insight into the genomic characteristics contributing to the MMR problem; (ii) the proportion of shared MMR between gene pairs provides information regarding the transcriptomic influences of mapping uncertainty for each dataset; and (iii) the degree of each gene, representing the number of gene-gene interactions resulting from (i) and (ii). GeneQC provides values for each extracted feature, a D-score, mapping uncertainty categorization, and alternative likelihood of categorization for each annotated gene corresponding to a provided dataset. More details of the procedure can be found in the following section.

If researchers continue processing RNA-Seq data with such high levels of mapping uncertainty (Figure 1A), all downstream analyses will have skewed and biased results. Just as raw reads require quality control—typically performed by FastQC (Andrews, 2010)—so do read alignment results. Even with tools that are specifically designed to address mapping uncertainty, such as *MMR* (Kahles, et al., 2015), the quality of the mapping results still requires investigation. Without some quality control for read alignment, researchers could potentially be using unreliable data, and blindly doing so.

## 2 Methods

### 2.1 GeneQC program and its implementation

GeneQC is designed to fit into computational pipelines for RNA-Seq data immediately following read alignment, acting as a supplement to most current pipelines. Currently, GeneQC is composed of two distinct processes: feature extraction and modeling. The feature extraction process is implemented using a Perl program. Modeling is performed on the feature extraction output using an R package, which provides the final output for GeneQC. More details on the implementation of GeneQC can be found in *Methods S1*.

### 2.2 Required Inputs for GeneQC

GeneQC takes as inputs three pieces of information that are easily found in most RNA-Seq analysis pipelines: (1) the read mapping result SAM file; (2) the fasta reference genome corresponding to that species; and (3) the species-specific annotation general feature format (gff/gff3) file (Figure 1B).

### 2.3 GeneQC Feature Extraction and Modeling

From input information, GeneQC performs feature extraction, in which the three characteristics are calculated for each annotated gene (Figure 1C): (i) maximum of the sequence similarity multiplied by the match length, where the match length is the longest continuous string of matching base pairs; (ii) maximum proportion of shared MMR between the gene of interest and another gene; and (iii) number of alternate gene locations with significant interactions with the gene of interest based on the previous two parameters. In addition to understanding the severity of the MMR problem in each sample, GeneQC provides species-or sample-specific insight into each feature’s impact on mapping uncertainty. This is done by developing a linear model to determine the significance and degree of impact for each feature. To perform the linear modeling, a dependent variable is constructed, with more detailed information on this value and each feature found in *Methods S2*.

GeneQC utilizes elastic-net regularization to develop a regression model for the calculation of D-scores. Elastic-net regularization is used to properly perform the variable selection, while simultaneously fitting a sufficient model to the provided data (Figure 1D). This approach also accounts for potential serious multicollinearity issues and prevents overfitting of the regression model. The set of calculated D-scores represents the mapping uncertainty for each annotated gene and is provided to give researchers an idea of how reliable their initial read mappings are, with a higher D-score representing more mapping uncertainty, and thus a less reliable expression estimate. More specific details regarding this process can be found in *Methods S3*.

Based on the calculated sets of D-scores through initial investigations during GeneQC development, there are apparent underlying distributions for these scores, intuitively representing levels of mapping uncertainty. For this purpose, extensive mixture model fitting is included within GeneQC to best fit a variable number of distributions to each set of D-scores (Figure 1E). Specifically, it is assumed that each set of D-scores can be expressed as a mixture model distribution given by

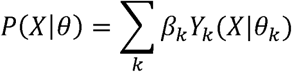

with *β*_*k*_ representing the weighting parameter of the *k*^*th*^ component, *Y*_*k*_ representing the probability density function of the *k*^*th*^ component of the mixture model, and *θ*_*k*_ representing the parameters of the *k*^*th*^ component. Based on our preliminary investigations into the D-score development, we have selected two underlying distributions for this purpose: Gamma and Gaussian. GeneQC fits mixture models for both the Gamma and Gaussian distributions with a variable number of distributions, ranging from two to five distributions. The optimally fitted mixture model is determined using a Bayesian Information Criterion (BIC) with a penalization based on the number of distributions is used to determine the best-fitting distribution. Specific details regarding the mixture model fitting procedure are outlined in *Methods S4*.

The best fitting mixture model is then used to separate each D-score into a category representing the severity of mapping uncertainty, thus indicating the mapping uncertainty categorization for each gene (Figure 1F). In addition to the mapping uncertainty categorization, an alternative likelihood value based on the posterior probabilities of the other distributions is provided to represent the certainty of the gene ID belonging to that category. Details for the categorization, cutoffs, and alternative likelihood are provided in *Methods S5*.

### 2.4 Generated Outputs from GeneQC

The final output of GeneQC includes the three extracted features, D-score, mapping uncertainty categorization, and alternative likelihood for each annotated gene. This information is combined into a concise table to provide users with all relevant information related to the mapping uncertainty of their read alignment data, along them to make informed decisions about further and continued analysis. An example output file is provided in *Methods S6* and *Table S2*.

## 3 Discussion and Conclusions

GeneQC is a tool used to investigate a prominent issue in modern RNA-Seq analysis. Oversight in the quality of RNA-Seq read mapping can have drastic consequences for all downstream analyses, and mapping uncertainty is a significant cause of problems in further analysis. While read mapping has been accepted as sufficient, entirely ignoring the possibility of poorly mapped reads used for further analysis can have detrimental effects on all manner of RNA-Seq studies. As demonstrated in our analysis of RNA-Seq data, the problem of mapping uncertainty is prominent and is displayed directly in the read alignment results. GeneQC can provide insight into the severity of this issue for each annotated gene based on genomic and transcriptomic features. It utilizes feature extraction, elastic-net regularization, and mixture model fitting to provide researchers with a sense of the quality of gene expression estimates resulting from the read alignment step. GeneQC provides sufficient information for researchers to make more well-informed decisions based on the results of their RNA-Seq data analysis and to plan further analyses to address this issue.

In addition to the direct provisions of GeneQC, interpretations of the coefficients allow for a further examination of the specific features contributing the mapping uncertainty. This will allow for further analysis and re-alignment strategies to be developed to the specific characteristics of the dataset. We are currently using this information to provide a computational tool capable of performing re-alignment of reads currently aligned to genes with high D-scores with the purpose of assisting researchers in the correction of mapping uncertainty. In the future, GeneQC will be integrated into a web server that applies this tool and associated re-alignment tools to perform large-scale RNA-Seq analyses on human, plant, and metagenome datasets. This application will allow for ease-of-use and collection of more data to support research with significant MMR issues.

## References

Anders, S. and Huber, W. Differential expression of RNA-Seq data at the gene level–the DESeq package. Heidelberg, Germany: European Molecular Biology Laboratory (EMBL) 2012.

Anders, S., Pyl, P.T. and Huber, W. HTSeq--a Python framework to work with high-throughput sequencing data. Bioinformatics 2015;31(2):166–169.

Andrews, S. FastQC: a quality control tool for high throughput sequence data. 2010.

Baruzzo, G., et al. Simulation-based comprehensive benchmarking of RNA-seq aligners. Nature methods 2017;14(2):135–139.

Bonfert, T., et al. ContextMap 2: fast and accurate context-based RNA-seq mapping. BMC bioinformatics 2015;16:122.

Chang, Z., et al. Bridger: a new framework for de novo transcriptome assembly using RNA-seq data. Genome biology 2015;16:30.

Chen, J., et al. ToppGene Suite for gene list enrichment analysis and candidate gene prioritization. Nucleic acids research 2009;37(suppl_2):W305–W311.

D’haeseleer, P. How does DNA sequence motif discovery work? Nature biotechnology 2006;24(8):959–961.

Dobin, A., et al. STAR: ultrafast universal RNA-seq aligner. Bioinformatics 2013;29(1):15–21.

Garber, M., et al. Computational methods for transcriptome annotation and quantification using RNA-seq. Nature methods 2011;8(6):469–477.

Ge, S.X. iDEP: An integrated web application for differential expression and pathway analysis. bioRxiv 2017.

Goff, L., Trapnell, C. and Kelley, D. cummeRbund: Analysis, exploration, manipulation, and visualization of Cufflinks high-throughput sequencing data. R package version 2013;2(0).

Grabherr, M.G., et al. Full-length transcriptome assembly from RNA-Seq data without a reference genome. Nature biotechnology 2011;29(7):644–652.

Harshbarger, J., Kratz, A. and Carninci, P. DEIVA: a web application for interactive visual analysis of differential gene expression profiles. BMC genomics 2017;18(1):47.

Kahles, A., Behr, J. and Rätsch, G. MMR: a tool for read multi-mapper resolution. Bioinformatics 2015;32(5):770–772.

Kim, D., Langmead, B. and Salzberg, S.L. HISAT: a fast spliced aligner with low memory requirements. Nature methods 2015;12(4):357–360.

Kong, Y. Btrim: a fast, lightweight adapter and quality trimming program for next-generation sequencing technologies. Genomics 2011;98(2):152–153.

Li, B. and Dewey, C.N. RSEM: accurate transcript quantification from RNA-Seq data with or without a reference genome. BMC bioinformatics 2011;12(1):323.

Li, B., et al. RNA-Seq gene expression estimation with read mapping uncertainty. Bioinformatics 2009;26(4):493–500.

Li, G., et al. QUBIC: a qualitative biclustering algorithm for analyses of gene expression data. Nucleic acids research 2009;37(15):e101–e101.

McDermaid, A., et al. ViDGER: An R package for integrative interpretation of differential gene expression results of RNA-seq data. bioRxiv 2018.

Miller, J.A., et al. Improving reliability and absolute quantification of human brain microarray data by filtering and scaling probes using RNA-Seq. BMC genomics 2014;15:154.

Nagalakshmi, U., et al. The transcriptional landscape of the yeast genome defined by RNA sequencing. Science 2008;320(5881):1344–1349.

Nelson, J.W., et al. The START App: a web-based RNAseq analysis and visualization resource. Bioinformatics 2017;33(3):447–449.

Nueda, M.J., et al. Identification and visualization of differential isoform expression in RNA-seq time series. Bioinformatics 2017.

Oshlack, A., Robinson, M.D. and Young, M.D. From RNA-seq reads to differential expression results. Genome biology 2010;11(12):220.

Ozsolak, F. and Milos, P.M. RNA sequencing: advances, challenges and opportunities. Nature reviews. Genetics 2011;12(2):87.

Pathan, M., et al. FunRich: An open access standalone functional enrichment and interaction network analysis tool. Proteomics 2015;15(15):2597–2601.

Perkel, J.M. Data visualization tools drive interactivity and reproducibility in online publishing. Nature 2018;554(7690):133–134.

Pertea, M., et al. Transcript-level expression analysis of RNA-seq experiments with HISAT, StringTie and Ballgown. Nature protocols 2016;11(9):1650–1667.

Pertea, M., et al. StringTie enables improved reconstruction of a transcriptome from RNA-seq reads. Nature biotechnology 2015;33(3):290–295.

Philippe, N., et al. CRAC: an integrated approach to the analysis of RNA-seq reads. Genome biology 2013;14(3):R30.

Pimentel, H., et al. Differential analysis of RNA-Seq incorporating quantification uncertainty. Nature methods 2017.

Powell, D. Degust: Visualize, explore and appreciate RNA-seq differential gene-expression data. In, COMBINE RNA-seq workshop. 2015.

Ritchie, M.E., et al. limma powers differential expression analyses for RNA-sequencing and microarray studies. Nucleic acids research 2015;43(7):e47.

Robinson, M.D., McCarthy, D.J. and Smyth, G.K. edgeR: a Bioconductor package for differential expression analysis of digital gene expression data. Bioinformatics 2010;26(1):139–140.

Subramanian, A., et al. Gene set enrichment analysis: a knowledge-based approach for interpreting genome-wide expression profiles. Proceedings of the National Academy of Sciences 2005;102(43):15545–15550.

Swan, M. The quantified self: Fundamental disruption in big data science and biological discovery. Big Data 2013;1(2):85–99.

Trapnell, C., et al. Differential analysis of gene regulation at transcript resolution with RNA-seq. Nature biotechnology 2013;31(1):46.

Trapnell, C., Pachter, L. and Salzberg, S.L. TopHat: discovering splice junctions with RNA-Seq. Bioinformatics 2009; 25(9):1105–1111.

Trapnell, C., et al. Differential gene and transcript expression analysis of RNA-seq experiments with TopHat and Cufflinks. Nature protocols 2012;7(3):562–578.

Wang, K., et al. MapSplice: accurate mapping of RNA-seq reads for splice junction discovery. Nucleic acids research 2010;38(18):e178.

Wang, Z., Gerstein, M. and Snyder, M. RNA-Seq: a revolutionary tool for transcriptomics. Nature reviews genetics 2009;10(1):57–63.

Wu, J., et al. OLego: fast and sensitive mapping of spliced mRNA-Seq reads using small seeds. Nucleic acids research 2013;41(10):5149–5163.

Wu, T.D., et al. GMAP and GSNAP for Genomic Sequence Alignment: Enhancements to Speed, Accuracy, and Functionality. Methods in molecular biology 2016;1418:283–334.

Wu, X., et al. Data mining with big data. IEEE transactions on knowledge and data engineering 2014;26(1):97–107.

Younesy, H., et al. VisRseq: R-based visual framework for analysis of sequencing data. BMC bioinformatics 2015;16(11):S2.

Yuan, L., et al. GAAP: Genome-organization-framework-Assisted Assembly Pipeline for prokaryotic genomes. BMC genomics 2017;18(Suppl 1):952.

Zhou, X. and Su, Z. EasyGO: Gene Ontology-based annotation and functional enrichment analysis tool for agronomical species. BMC genomics 2007;8(1):246.

